# Two-fold Red Excess (TREx): A simple and novel digital color index that enables non-invasive real-time health assessment of green-leaved as well as anthocyanin-rich crops

**DOI:** 10.1101/2024.12.08.627401

**Authors:** Avinash Agarwal, Filipe de Jesus Colwell, Viviana Andrea Correa Galvis, Tom R. Hill, Neil Boonham, Ankush Prashar

## Abstract

**Background:** Digital color indices provide a reliable means for assessing plant health status by enabling real-time estimation of chlorophyll (Chl) content, and are thus adopted widely for crop monitoring. However, as all prevalent leaf color indices used for this purpose have been developed using green-leaved plants, they do not perform reliably for anthocyanin (Anth)-rich red-leaved varieties. Hence, the present study investigates digital color indices for six types of leafy vegetables with different levels of Anth to identify congruent trends that could be implemented universally for non-invasive crop monitoring irrespective of species and leaf Anth content. For this, datasets from three digital color spaces, viz., RGB (Red, Green, Blue), HSV (Hue, Saturation, Value), and *L*a*b\** (Lightness, Redness-greenness, Yellowness-blueness), as well as various derived plant color indices were compared with SPAD Chl meter readings and Anth/Chl ratio of *n* = 320 leaf samples.

**Results:** While most digital color features and indices presented abrupt shifts between Anth-rich and green-leaved samples, the newly-developed color index Two-fold Red Excess (TREx) as well as the color feature R showed very strong correlation with SPAD readings (*R*^2^ > 0.84), and did not exhibit any deviation due to leaf Anth content. Moreover, both parameters could predict SPAD values reliably (*R*^2^ > 0.75). Further, logarithmic decline of G/R and Augmented Green-Red Index (AGRI) with increasing Anth/Chl ratio (*R*^2^ > 0.82) revealed that relative Anth content affected digital color indices markedly by shifting the R:G balance until the Anth/Chl ratio reached a certain threshold.

**Conclusion:** The present study provides the first in-depth assessment of variations in RGB-based digital color indices due to high leaf Anth contents, and uses the data for Anth-rich as well as green-leaved crops belonging to different species to develop a universal digital color index TREx that can be used as a reliable alternative to handheld Chl meters for rapid high-throughput monitoring of green-leaved as well as red-leaved crop varieties.

## 1. Background

Transition of emphasis from “food quantity” to “food quality” has led to a noticeable surge in interest towards expanding and improving the production of leafy vegetables in the past decade (1). Growing focus on the nutritive value of foodstuffs has highlighted the importance of anthocyanins (Anth) as key nutritional compounds owing to their high antioxidant activity and numerous potential health benefits (2). Consequently, there have been concerted efforts to promote large-scale production of various Anth-rich leafy vegetables belonging to diverse plant families (3). This rising interest in cultivation of Anth-rich “red-leaved” vegetables has brought to light a new challenge for growers: monitoring the health status of such crops efficiently.

Leaf chlorophyll (Chl) content has been widely used an indicator of plant health status since decades as it is connected strongly with plant nitrogen content and directly affects photosynthesis (4–6). Hence, non-invasive monitoring of crops via hand-held Chl meters has become very common in the past decades (7–10). However, such devices require manual measurements from each leaf, making the process slow and labor-intensive. Further, inferences from point measurements are subjective considering localized variations in pigmentation within the leaf. As an alternative, various machine vision technologies have emerged as reliable means of high-throughput real-time monitoring of plants (11,12). Amongst those, RGB (Red, Green, Blue) cameras have been adopted most widely for crop monitoring considering the synergistic relation between plant health status, Chl content, and leaf color (9,13–15). Steady improvements in RGB sensors have resulted in better resolution, reduced size, easy availability, and hassle-free application of RGB cameras, making it highly feasible for crop monitoring at a commercial scale (16–18). Hence, RGB imaging has been especially well explored for studying the variations in leaf color which “reflect” the physiological status of plants.

However, because green-leaved plants dominate conventional commercial cultivation, the existing digital color indices for crop monitoring have been developed primarily for plants with low Anth levels. Consequently, such indices focus on the total and relative abundance of Chl and carotenoids (Car) as the key indicators of plant health status (9,13–15,19–22). Conversely, digital image analysis for red-leaved (Anth-rich) plants has primarily focused on estimating leaf Anth content (23–25). A few studies have demonstrated the feasibility of predicting Chl content in sweet potato (26) and various Anth-rich tree leaves (27,28) using reflectance spectra. However, implementation of RGB-based color indices for monitoring green-leaved as well as Anth-rich plants in tandem remains largely unexplored.

Therefore, the current study aims to develop a universal method for monitoring both green-leaved and Anth-rich plants using RGB imaging. For this, we analyze images from six different leafy vegetables with varying levels of Chl and Anth to visualize the impact of high Anth content on digital color features as well as established and newly-developed color indices. The investigation focuses on identifying digital color attributes that remain consistent despite variations in Anth content, and hence, could be implemented in non-invasive real-time crop monitoring across different species and leaf Anth levels. Additionally, the investigation also explores the impact of Anth/Chl ratio on digital color parameters providing insights into how different pigment combinations influence leaf color profiles.

## 2. Materials and methods

### 2.1. Plant material

Six leafy vegetables with different levels of Anth content were selected for the present study (Fig. 1), and were classified into three groups based on leaf pigment status as follows: 1) high Anth (HA) – Purple basil (*Ocimum basilicum* L. var. *purpurascens*; PB) and Red pak choi (*Brassica rapa* L. ssp. *chinensis* cv. ‘Rubi F1’; RPC); 2) medium Anth (MA) – Scarlet kale (*Brassica oleracea* L. var. *acephala ‘Scarlet’*; SK); and 3) low Anth (LA) – Green pak choi (*Brassica rapa* L. ssp. *chinensis*; GPC), arugula (Eruca vesicaria ssp. sativa Mill. cv. ‘Wasabi Rocket’; WR), and Greek basil (*Ocimum basilicum* L. var. *minimum*; GB). Leaf color ranged between dark purple and reddish-green for HA samples, green lamina with reddish-tinge and red midrib for MA samples, and different yellow-green shades with no hint of red for LA samples (Fig. 1).

**Fig. 1.**
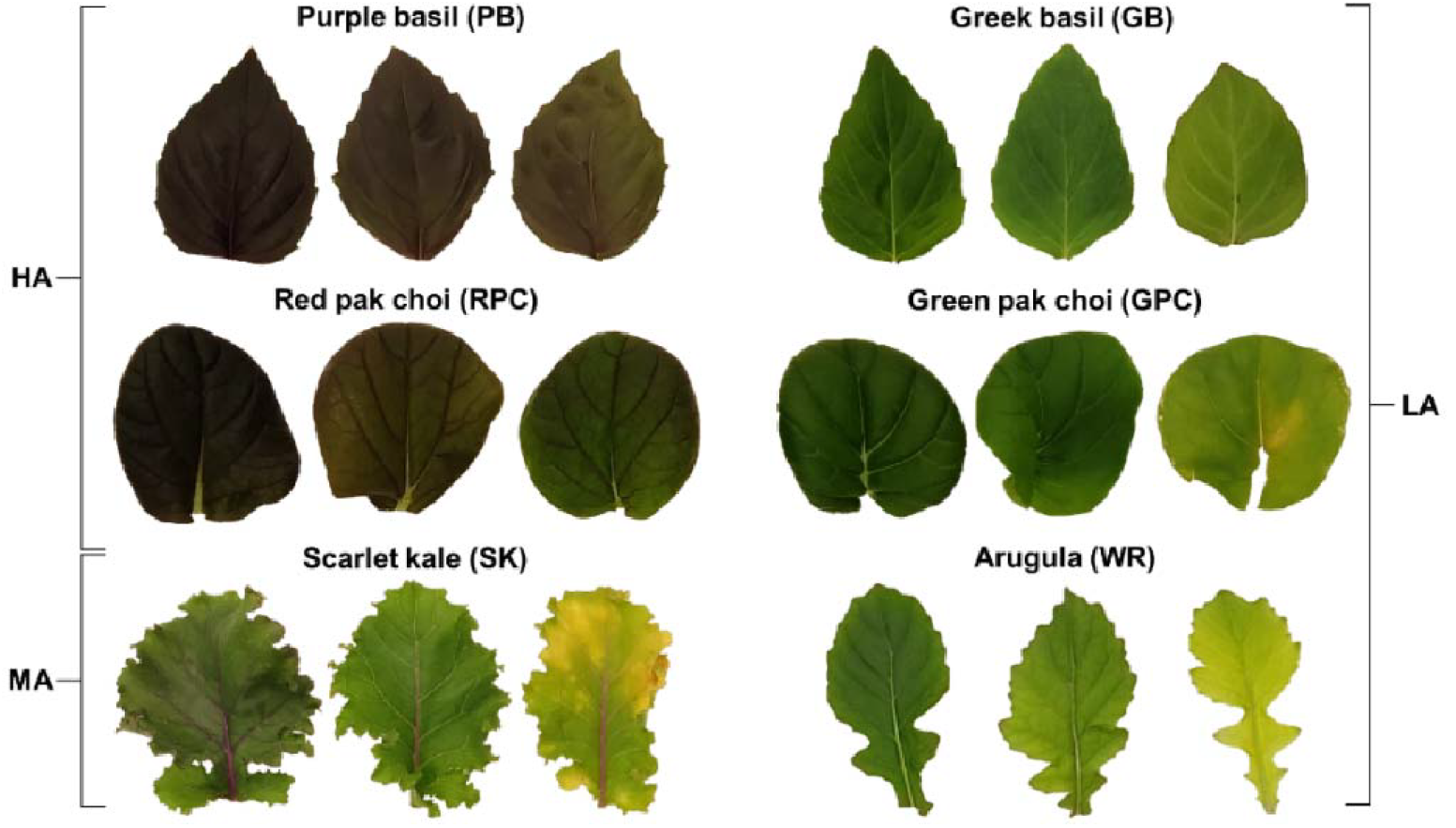
Representative leaf samples with varying levels of pigmentation from the six types of leafy vegetables selected for the present study. Purple basil and Red pak choi were grouped as high anthocyanin (HA) samples, and Scarlet kale was considered as the only member of the medium anthocyanin (MA) group. Greek basil, Green pak choi, and Arugula (cv. ‘Wasabi rocket’; WR) were categorized as low anthocyanin (LA).

Seedlings of all plants were grown in coco-peat plugs (Van der Knapp, The Netherlands) in a nursery (Aralab-InFarm UK Ltd., London, UK), with each plug holding 5–10 seedlings. When the seedlings reached a height of *ca*. 5 cm, the seedling plugs were transferred to an experimental hydroponic vertical farm (InStore Farm V2, InFarm UK Ltd.) stationed at the Agriculture Building, Newcastle University, UK. Seedling plugs for each type of plant were placed in two hydroponic trays (30×40 cm^2^) having a 3×4 array of equally-spaced empty slots for seedling plugs, totaling 24 seedling plugs for each plant type. A commercial hydroponics fertilizer mix was used as the nutrient source, and irrigation was performed following the ebb-and-flow system wherein the nutrient solution was flooded into the hydroponic chamber intermittently (10 min/h) to soak the roots. A white LED array having an approximate red (400–499 nm):green (500–599 nm):blue (600–699 nm) distribution of 40:20:40 was used to provide a PPFD of 280 µmol/m^2^ sec following a 16/8 h day-night cycle. Temperature and relative humidity were maintained at 25±1 °C and 65±5%, respectively. Sensors for temperature, humidity, flow rate, electrical conductivity, and pH within the vertical farming system were connected to a Farmboard (InFarm UK Ltd.) for real-time monitoring of the plant growth environment.

### 2.2. Leaf sampling and SPAD measurement

A total of *n* = 320 leaf samples were collected from the six types of plants (PB, *n* = 60; RPC, *n* = 40; SK, *n* = 100; GPC, *n* = 40; WR, *n* = 40; GB, *n* = 40) between 15–20 days of growth within the experimental setup. Samples with variations in pigmentation due to inherent physiological changes were selected to get a wide range of leaf color profiles, whereas very young (<10 days old) as well as fully-senesced leaves were specifically avoided. Leaves were labeled prior to excision, and Chl content was measured non-invasively by taking three readings from each leaf via a SPAD-502 Plus meter (Konica-Minolta, Inc., Tokyo, Japan) avoiding the midrib and prominent veins (29). SPAD measurements have been considered the “gold standard” for non-invasive assessment of plant health status and estimation of leaf Chl content following numerous reports over more than two decades (10), and hence, have been used likewise in the present study. Subsequently, leaves were excised at the base for image acquisition (described in the next section), followed by destructive measurement of pigment contents (described later).

### 2.3. Image acquisition

Leaf samples were transferred to a customized imaging setup for digital image acquisition (Fig. 2) immediately following excision. The setup comprised of a metal frame for mounting a smartphone camera and LED-luminaires for lighting, along with a horizontal platform (stage) with a matte white surface for placing the leaf samples. Images were acquired using a Redmi Note 7 Pro smartphone (Xiaomi Corp., Beijing, China) equipped with a Sony IMX 586 RGB sensor (size 1/2.0", Quad-Bayer array) within a dual rear-camera system (primary lens: resolution 48 megapixels, aperture *f*/1.8, wide angle, pixel size 1.6 µm, phase detection autofocus; secondary lens: resolution 5 megapixels, aperture *f*/2.4, depth perception). The Open Camera android application (ver. 1.52, developer: Mark Harman, source: Google Play Store) was used for capturing images (8000×6000 pixels, sRGB color space, JPEG format). The smartphone was placed within a compact cradle suspended from the metal frame to prevent camera movement or change in camera angle.

**Fig. 2.**
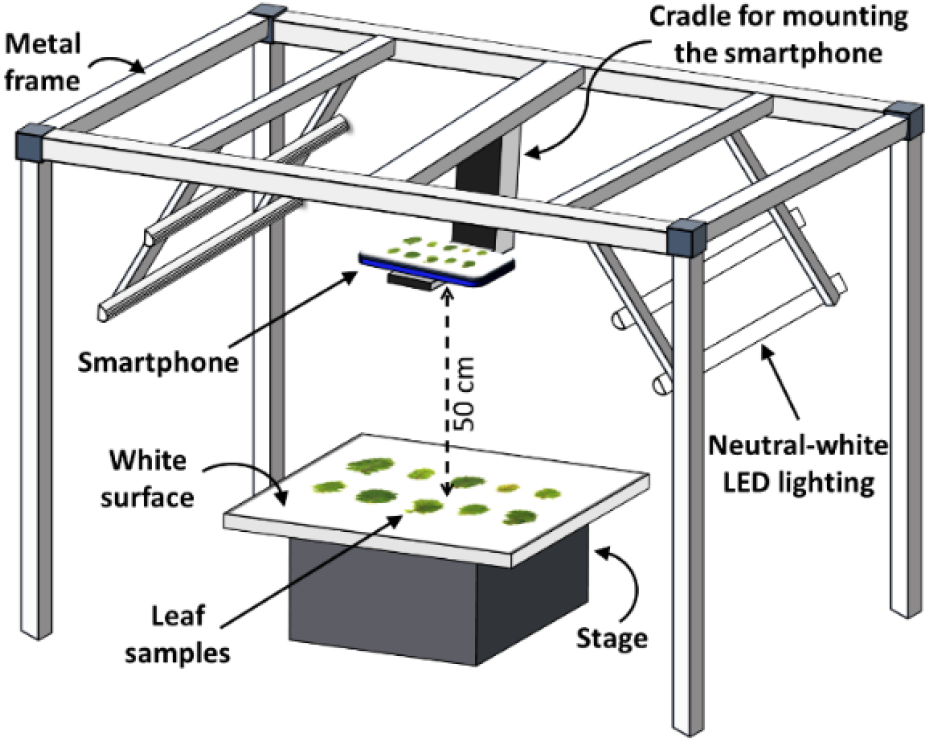
Schematic representation of the customized setup for leaf image acquisition. A metal frame was used for mounting a smartphone camera and LED lights. A matte white board was used as the background for leaf imaging while maintaining a fixed distance of 50 cm from the camera. Camera parameters (focus, exposure, and ISO) were set by focusing on the empty stage to maintain uniformity of color tone across images. Images were captured using the voice-activated mode to operate the camera remotely, avoiding camera movement and shadows.

Camera-to-stage distance of 50 cm was maintained along with constant imaging parameters (exposure time 1/100 sec, ISO-200). Camera focus was fixed on the stage before placing the leaf samples, and automatic adjustments (autofocus and exposure compensation) were disabled to ensure uniformity across images.

Camera flash was disabled during the process to avoid glares. Instead, lighting was provided by four neutral-white (4000 K) LED tube-lights (Model No. 0051048, Feilo Sylvania International Group Kft., Budapest, Hungary; www.sylvania-lighting.com). Fixed lighting prevented undesirable fluctuations in brightness and color temperature between images, and nullified image pre-processing requirements. As enough lighting was provided, a relatively low ISO was adequate for ensuring sharp images while minimizing noise. Images were captured remotely using the voice-activated mode within the smartphone application to avoid shadows, delays, or any disturbances that could be caused by manual handling.

### 2.4. Spectrophotometric estimation of pigment contents

Leaf sections were collected for spectrophotometric estimation of total Chl, Car, and Anth contents immediately following image acquisition. Briefly, two 2 cm^2^ sections were excised from each leaf, weighed individually, sealed into separate 1.5 ml centrifuge tubes, and transferred to −20 °C for storage. Vials of all the samples were subsequently put in a liquid nitrogen bath, followed by tissue pulverization using stainless-steel beads within a tissue homogenizer (Geno/Grinder 2010, SPEX SamplePrep, Cole-Parmer, Illinois, USA). One batch of samples (*n* = 320) was used for estimating Chl and Car contents, whereas the other batch (*n* = 320) was used for estimating Anth content.

Chl and Car contents were estimated following acetone extraction as described by Lichtenthaler (30). Briefly, 1 ml of ice-cold 80% (v/v) acetone was added to each vial, followed by centrifugation at 10,000 g at 4 °C for 15 min. The supernatant was collected, and the pellet was re-extracted using 1 ml of the solvent. Both supernatants were pooled, and absorbance was recorded spectrophotometrically at 470 nm (A_470_), 647 nm (A_647_), and 663 nm (A_663_) for calculating the total Chl and Car contents per unit leaf fresh weight (FW) for 2 ml (Vol) of extract as follows:

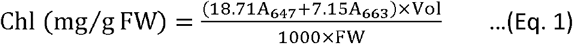

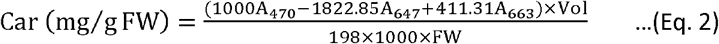

Anth content was estimated following the method of Mancinelli et al. (31). A similar extraction procedure as above was followed using chilled acidified (1% w/v HCl) methanol as the solvent. Absorbance was recorded at 530 nm (A_530_) and 657 nm (A_657_) for calculating the total Anth content using the expression A_530_ – (0.25×A_657_), where A_530_ corresponds to the peak absorbance of Anth and A_657_ was used for pheophytin correction. A standard curve was prepared using cyanidin-3-O-glucoside (Merck KGaA, Darmstadt, Germany) for calculating relative Anth content per unit leaf biomass (mg/g FW).

### 2.5. Color feature extraction

A customized image processing pipeline was designed to extract the color feature values of whole leaves using the *numpy* and *cv2* libraries in Python program (www.python.org). Within the pipeline, each RGB image was used to directly extract the R, G, B features using *cv2* library commands. Additionally, Hue, Saturation, and Value (HSV) as well as Lightness, Redness-greenness, and Yellowness-blueness (*L*a*b\**) color features were also extracted from the image. For each color feature channel, values for all pixels within the leaf boundary (minimum 5000 pixels) were averaged to obtain the mean color feature value for the entire leaf. Normalized RGB features and various previously-reported leaf color indices (Table 1) were calculated using these color feature values. In addition, we introduced a new redness-based color index, Two-fold Red Excess (TREx; Table 1).

**Table 1.**
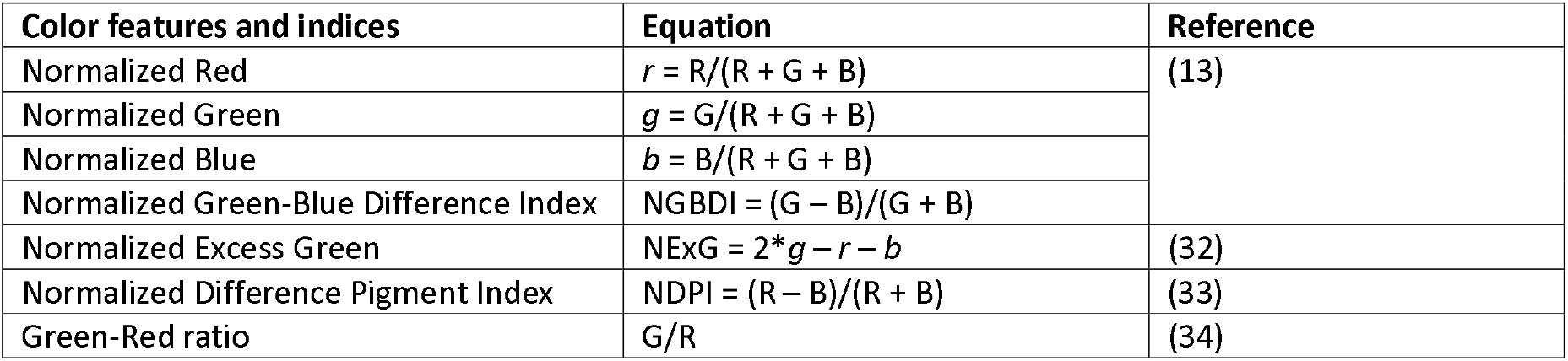
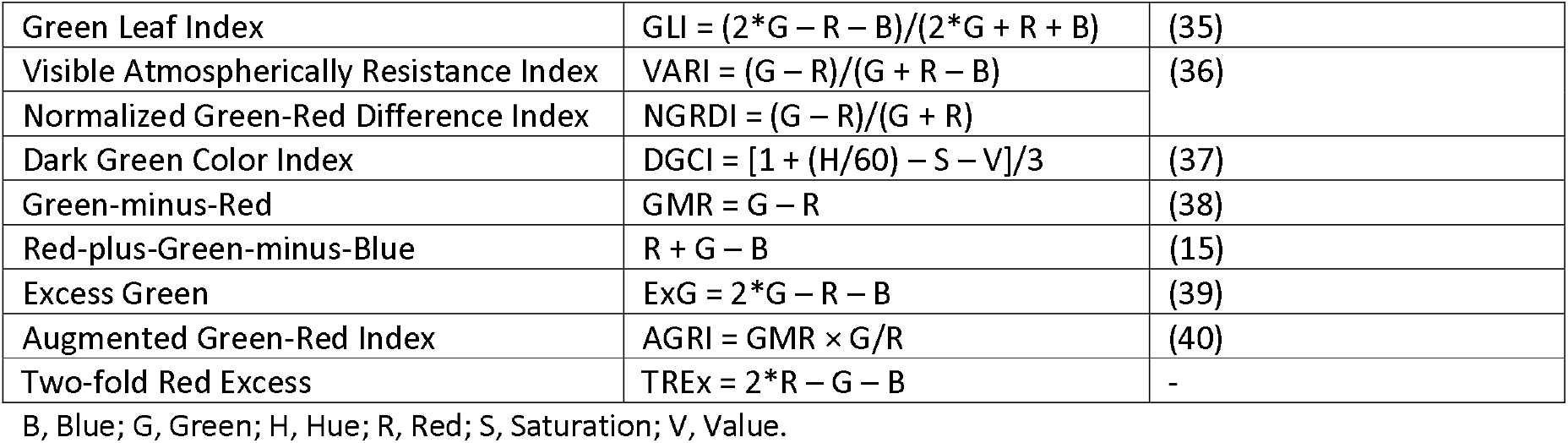
Normalized RGB features and vegetation indices derived from digital images.

### 2.6. Data visualization and comparison

Averaged SPAD values (*n* = 3 per leaf) were plotted against the spectrophotometrically evaluated Chl content by collating the data for HA, MA, and LA samples to visualize relative trends. Similarly, Car and Anth contents were also plotted against Chl content to compare relative trends across the different sample categories. Subsequently, RGB, HSV, *L*a*b\**, and *rgb* values were individually compared with the SPAD values as well as Anth content to visualize the variations in different digital color features across the three categories. Further, selected vegetation indices were plotted against SPAD values to visualize the trends for HA, MA, and LA samples. Similarly, Anth/Chl ratio was compared with individual color features as well as selected color indices to understand the impact of different pigment blends on RGB image-based color attributes.

### 2.7. SPAD value prediction

Digital color features and indices showing good correlation with SPAD readings (*R*^2^ > 0.7) were used to predict SPAD values using the entire dataset (*n* = 320) as well as following five-fold cross validation (Training: *n* = 256; Test: *n* = 64) within R-software using the ggplot2 library. Linear, quadratic, and logarithmic curve-fitting was tested using the *“lm”* function, followed by SPAD value prediction using the *“predict”* function. The *‘‘summary’’* function was used to obtain *R*^2^, root-mean-squared error (RMSE), level of significance (*p*), and coefficient values for each model. Equations for models with the best performance were chosen.

### 2.8. Statistical analysis

Homogeneity of variance for Chl, Anth, and Car contents across the different plant species and varieties was assessed using the Bartlett test. Subsequently, Kruskal-Wallis non-parametric test and Dunn’s post hoc test were performed to assess the significance of difference amongst the different leafy vegetables for the content of each type of pigment. Curve-fitting via linear, quadratic, exponential, logarithmic, and power functions was performed in Microsoft Excel 365 (Microsoft Corp., USA) for obtaining regression trends for all scatter plots (*n* = 320). Equations along with the respective coefficient of determination (*R*^2^) for best-fit trendlines were selected to represent the relation mathematically. Point of inflection (elbow) for the best-fit curve of color indices in relation to Anth/Chl ratio was detected in R-software (ver. 4.0.3; www.rproject.org) within the R-Studio environment (ver. 1.3.1093; www.rstudio.com) using the inflection package following the Extremum Distance Estimator method.

## 3. Results

### 3.1. Pigment contents and SPAD readings

Chl as well as Car contents were comparable for all types of leafy vegetables, whereas Anth content varied significantly as expected (Fig. 3a). Specifically, Anth content was highest (*p* < 0.05) in the HA plants, i.e., PB followed by RPC. Further, while mean Anth content for the MA group, i.e., the SK samples, was relatively closer to the lower range, it was lower still in the three LA plants, i.e., GB, WR, and GPC. A strong positive non-linear correlation was observed between Chl content and SPAD readings (*R*^2^ = 0.804; *n* = 320) upon combining the data for all samples (Fig. 3b). In contrast, the comparison between Chl and Car contents revealed a strong positive linear relation (*R*^2^ = 0.705; *n* = 320; Fig. 3c). However, no clear trend was evident between Anth and Chl contents (*R*^2^ < 0.1; *n* = 320), indicating a lack of correlation between the contents of these two pigments (Fig. 3d).

**Fig. 3.**
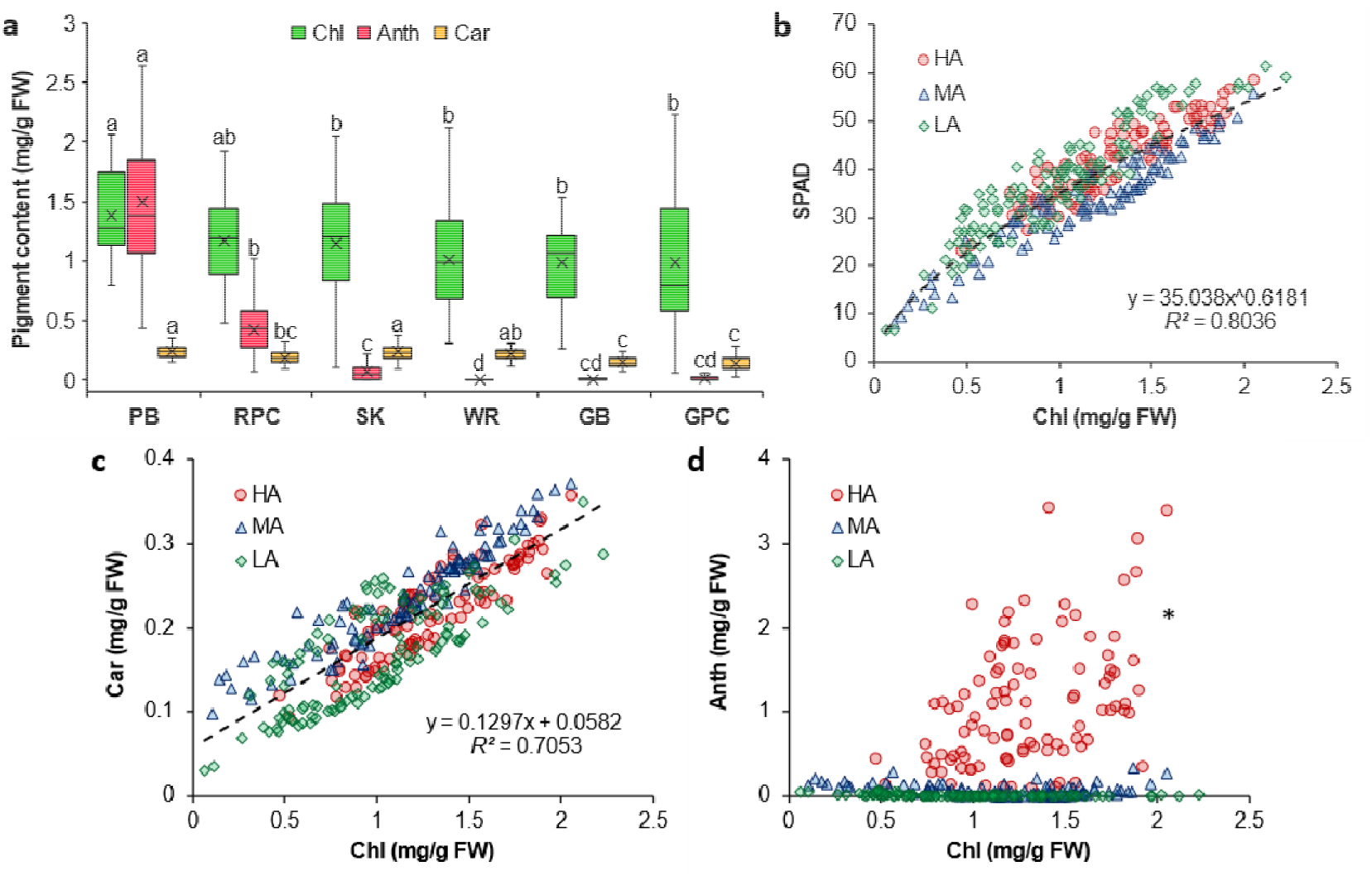
Chlorophyll (Chl), anthocyanin (Anth), and carotenoid (Car) contents of Purple basil (PB; *n* = 60), Red pak choi (RPC; *n* = 40), Scarlet kale (SK; *n* = 100), Arugula cv. ‘Wasabi rocket’ (WR; *n* = 40), Greek basil (GB; *n* = 40), and Green pak choi (GPC; *n* = 40) (a), as well as the relation of Chl content with SPAD values (b), Car content (c), and Anth content (d) for plants with high (HA), medium (MA), and low (LA) levels of Anth. Box-and-Whisker plots show the mean (×), median (horizontal line), interquartile range (box), and whiskers representing 5 and 95% percentiles (a). Significant differences in mean values for each type of pigment (a) are indicated by different alphabets as per Dunn’s post hoc test (*p* < 0.05). Equations (b, c) describing the best fit curves for all data combined (*n* = 320) have been presented along with the coefficients of determination (*R*^2^). *Fitted curve for Chl vs Anth (d) has not been presented owing to very poor correlation (*R*^2^ < 0.1; *n* = 320). FW, fresh weight.

### 3.2. Comparing color indices with SPAD readings

All digital color indices revealed characteristic trends upon plotting with SPAD values (Fig. 4). In general, HA samples clustered distinctly from the MA and LA samples for most of the indices. This highlighted the misleading effect of high Anth content on the digital color information of red-leaved plants. Similar results were obtained upon plotting SPAD readings and Anth content against most color features (Supplementary Figs. S1, S2). As an exception, plots of SPAD against the TREx color index (Fig. 4i) and R color feature (Supplementary Figs. S1a) did not show any deviation due to the presence of Anth, and a homogenous distribution of data points was observed for both parameters upon collating the information for HA, MA, and LA samples (0.845 < *R*^2^ < 0.855; *n* = 320). DGCI also exhibited homogenous data distribution (Fig. 4e), but had relatively weaker correlation with SPAD (*R*^2^ = 0.71; *n* = 320) compared to TREx and R.

**Fig. 4.**
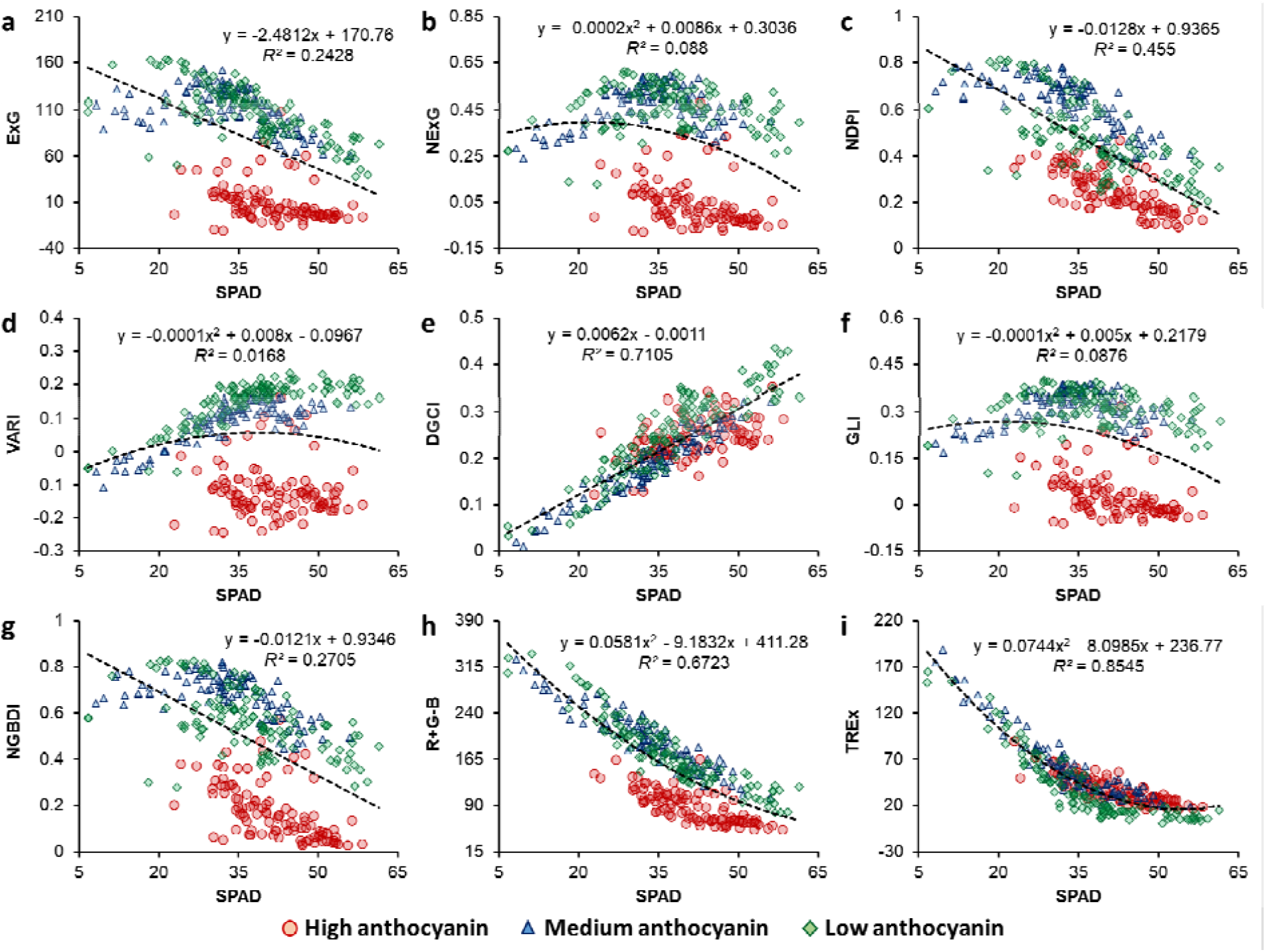
Plots for SPAD measurements versus Excess Green index (ExG; a), Normalized Excess Green index (NExG; b), Normalized Difference Pigment Index (NDPI; c), Visible Atmospherically Resistance Index (VARI; d), Dark Green Color Index (DGCI; e), Green Leaf Index (GLI; f), Normalized Green-Blue Difference Index (NGBDI; g), Red-plus-Green-minus-Blue index (R+G-B; h), and Two-fold Red Excess index (TREx; i) for leafy vegetables with different levels of anthocyanin (indicated with different symbols). Coefficients of determination (*R*^2^) and equations have been presented for the best-fit curve for the combined dataset (*n* = 320).

### 3.3. Comparing color indices with pigment ratios

Anth/Chl ratio ranged between 0.05–2.5 for the HA category, whereas it was generally less than 1.5 for the MA group and below 0.05 for the LA samples, with a few exceptions due to extremely low Chl in some LA leaves (Fig. 5). Amongst all color indices, G/R and AGRI showed the best correlation with Anth/Chl ratio (*R*^2^ > 0.82; *n* = 320), wherein samples from all three groups were clustered homogenously, irrespective of leaf Anth status. Color indices NGRDI, GMR, VARI, and GLI also exhibited similar results, albeit with slightly poorer correlation (0.68 < *R*^2^ < 0.82; *n* = 320). Point of inflection (elbow) was found to be at Anth/Chl ≈ 0.2 for all six indices (Fig. 5). The other indices (data not shown) generally resulted in the segregation of HA, MA, or LA samples based on Anth/Chl ratios or did not show any clear trend (*R*^2^ < 0.1; *n* = 320), and hence, were not considered for further discussion. Amongst individual color features, H showed good correlation with Anth/Chl values (*R*^2^ = 0.816; *n* = 320), followed by *a\** (*R*^2^ = 0.746; *n* = 320), while the other features did not yield any reliable trends (Supplementary Fig. S3).

**Fig. 5.**
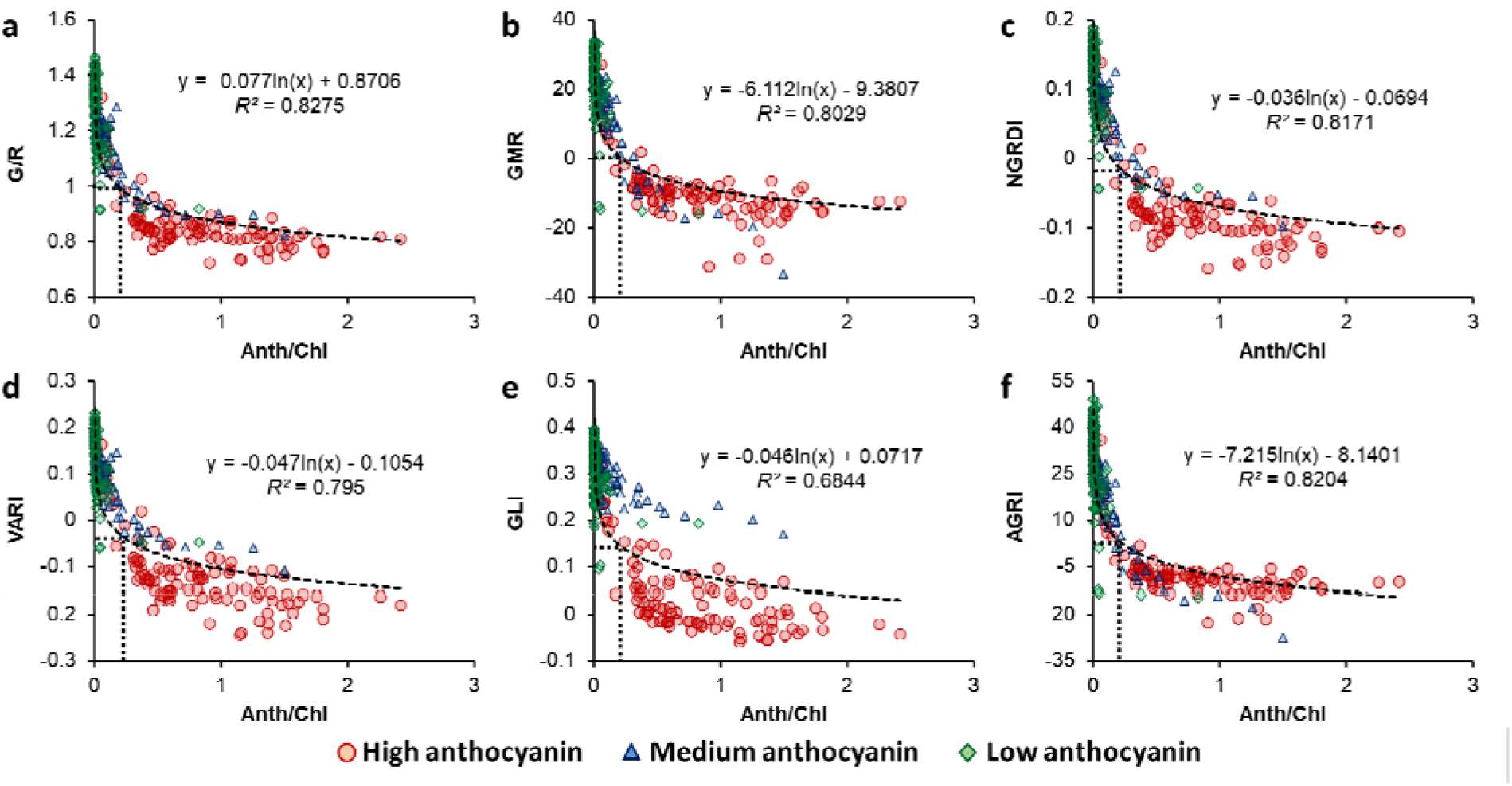
Plots for anthocyanin/chlorophyll ratio (Anth/Chl) versus Green/Red ratio (G/R; a), Green-minus-Red index (GMR; b), Normalized Green-Red Difference Index (NGRDI; c), Visible Atmospherically Resistance Index (VARI; d), Green Leaf Index (GLI; e), and Augmented Green-Red Index (AGRI; f) for leafy vegetables with different levels of anthocyanin (indicated with different symbols). Coefficients of determination (*R*^2^) and equations have been presented for the best-fit curve of the combined dataset (*n* = 320). Dotted rectangle indicates the point of inflection (elbow) in the curve.

### 3.4 Predicting SPAD values

Prediction of SPAD values was attempted using R, TREx, and DGCI following good correlation with SPAD measurements (*R*^2^ > 0.7) and homogenous distribution of data for HA, MA, and LA samples (Fig. 4e, 4i, Supplementary Fig. S1a). Amongst the three digital color attributes, R predicted SPAD values most accurately (*R*^2^ = 0.791, RMSE = 4.735), followed by TREx (R = 0.753, RMSE = 5.152) and DGCI (*R*^2^ = 0.711, RMSE = 5.573) as depicted (Fig. 6). Results of five-fold cross-validation yielded similar results (Supplementary Tables S1–S3), indicating consistency of predictive performance for all three parameters with unseen data as well as with different training datasets.

**Fig. 6.**
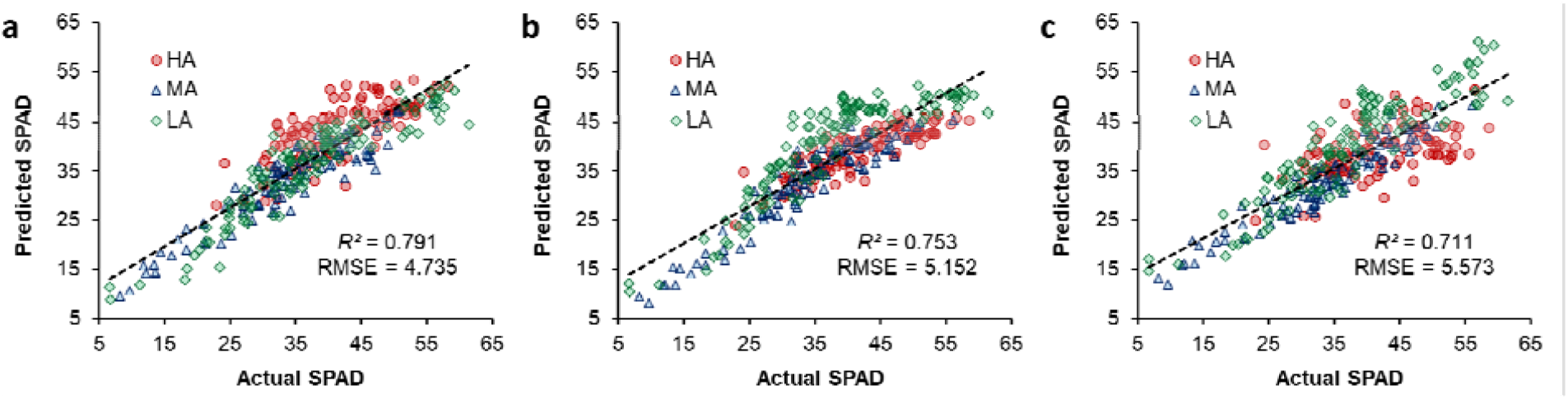
Actual versus predicted SPAD values obtained using R (a), TREx (b), and DGCI (c) values of leaves with high (HA), medium (MA), and low (LA) anthocyanin contents. Coefficient of determination (*R*^2^) and root-mean-squared error (RMSE) have been shown for the models generated by collating the data for all three groups (*n* = 320; *p* < 0.001). Equations of prediction models and results of five-fold cross-validation are provided as supplementary tables.

#### 4. Discussion

### 4.1. SPAD readings and digital color trends for varying pigment blends

Leaf pigment composition and cellular organization determines the absorption, transmission, and reflectance of incident light (16). Since each type of leaf pigment captures photons from specific regions of incident light, low proportions of the respective photons are present in light reflected from the leaf surface. Hence, the perceived leaf color represents the pigment that is capable of photon absorption within that waveband, but is present at a low concentration (41). For instance, leaves with adequate Chl content appear green when the concentration of green-light-absorbing pigment such as Anth is relatively low. Similarly, an Anth-rich leaf appears red only when the pigment primarily responsible for the absorption of red light, i.e., Chl, is present at low concentrations (42–44). Accordingly, leaves with very high contents of both Chl and Anth appear dark-purplish due to strong absorption of photons across the entire visible spectrum (45,46), as also seen in this study for the HA samples (Fig. 1). Conversely, a leaf with very low concentrations of both Chl and Anth appears “yellowish” because of high reflectance in both red and green wavebands, which indicates unmasking of Car (47).

In this context, selection of leaf samples from six different plant species and varieties displaying a wide range of visual color profiles (Fig. 1) enabled a comprehensive investigation of all possible distinctions in digital color attributes due to variations in leaf pigment blends (Fig. 3a). The strong correlation of SPAD measurements with spectrophotometrically measured Chl contents (*R*^2^ = 0.804; *n* = 320; Fig. 3b) reiterated the versatility of the SPAD meter as a reliable indicator of leaf Chl status and plant health (10,15,48,49), and also highlighted its uniformity across multiple plant species (7–9,29). Notably, there was no impact of Anth content on SPAD values, which is understandable considering that SPAD meters measure photon transmission only at 650 and 940 nm (34), i.e., within the red and infrared wavebands wherein Anth pigments are not very active.

Poor correlation between Chl and Anth contents (*R*^2^ < 0.1; *n* = 320; Fig. 3d) was not only expected, but was also essential for the current investigation as it provided a wider palette of leaf colors due to different pigment blends allowing more in-depth assessment of changes in digital color attributes. In contrast, the positive linear correlation between Chl and Car contents across all six types of leafy vegetables (*R*^2^ = 0.705; *n* = 320; Fig. 3c) indicated a strong synergy between the contents of both these pigments, with similar findings having been reported in other plant species as well (33,47,50,51). In green-leaved plants, increase in Car/Chl ratio with decreasing Chl content causes leaf yellowing due to unmasking of Car pigments, acting as an indicator of leaf senescence or declining health (50,52,53), and this phenomenon forms the basis of commonly-used plant color indices. However, the influence of Car was not very clear in this study considering its markedly lower concentration relative to the other pigments in most of the samples (Fig. 3). Hence, the present study focused on Anth/Chl instead of Car/Chl to better understand the dynamics of leaf color variations caused by high Anth content.

Anth/Chl ratio was found to be crucial in determining leaf color profile. Specifically, increase in the relative abundance of Anth, i.e., increasing Anth/Chl ratio, caused a rapid decline in G/R and GMR values till G and R values converged, i.e., *ca*. G/R = 1 and GMR = 0 was reached (Fig. 5a, b). This rapid decline in index values was also evident for NGRDI, VARI, GLI, and AGRI as well (Fig. 5c–f). Notably, the rate of change in all six color indices decreased considerably till the Anth/Chl ratio of about 0.2 (Fig. 5), with further increases in Anth/Chl ratio resulting in relatively nominal change in the respective indices. This is indicative of a threshold effect of Anth content on color indices, i.e., increasing Anth/Chl ratio influences color indices only till a certain level but has limited effect thereafter. Comparing Anth/Chl ratio with different color features revealed that the R and G channels did not correlate well with different pigment blends individually (Supplementary Fig. S3a, b). Instead, features such as H and *a\**, which inherently account for the balance between redness and greenness, were more effective in accurately representing the relative abundance of Anth and Chl (Supplementary Fig. S3d, h). These observations clearly indicate that increase in leaf Anth alters the balance between R and G features considerably, which results in significant changes in color index values until Anth/Chl ratio reaches a certain threshold. This also explains why most of the conventional color indices which rely on the R:G balance of green-leaved plants gave variable results for Anth-rich samples.

### 4.2 Selecting the color index for broad-spectrum crop monitoring

As Chl content is a strong indicator of plant health, identification of digital color indices for crop monitoring has always focused on finding correlations with leaf Chl content estimated via leaf extracts and/or Chl-meter measurements (49,54–56). Indices such as DGCI (15,21), NGRDI, VARI (49), ExG (55), and NDPI (57), as well as R and G color features (14,15,19,58) have been shown to correlate well with extract-based as well as non-invasive Chl content estimates. Notably, all such studies have focused on green-leaved crops, such as soybean, potato, spinach, coffee, barley, and wheat, with hardly any reports on crops with anthocyanic leaves.

In the present study, comparison of SPAD values with these well-established color indices yielded unexpected variations in data distribution and correlation upon using the information for HA, MA, and LA samples. For example, ExG = 50, corresponding to a SPAD value of *ca.* 55 for green-leaved (LA) samples, was found to coincide with SPAD readings of ca. 30 for Anth-rich (HA) samples (Fig. 3a). This would imply that ExG of a healthy green-leaved plant would be the same as that of a relatively unhealthy Anth-rich plant, signifying the misleading or “red herring” effect of leaf Anth in RGB analyses. Similar trends were observed for indices such as NDPI, NGBDI, and R+G-B (Fig. 3c, g, h). On the other hand, indices such as NExG, VARI, and GLI showed a bifurcation of the dataset, with LA and MA samples clustered together, but having negligible overlap with HA samples along the ordinate of the respective indices despite similar SPAD values (Fig. 3b, d, f). In contrast, DGCI, a vegetation index based on the HSV color space, did not differentiate between HA and the other two groups, and provided a better overall correlation with SPAD values (*R*^2^ = 0.71; *n* = 320; Fig. 4e) compared to the other established color indices. While the availability of additional HSV-based color indices would provide more insights into the impact of leaf Anth content within this color space, observations for the other indices clearly indicate that there is a major shift in the RGB color space data which limits the scope of implementing existing RGB-based indices for monitoring green-leaved and Anth-rich crops in tandem.

Since the color “green” is intuitively associated with leaves, the existing digital color indices formulated for assessing plant health have often focused on evaluating the “greenness” of leaves (32,35,37,39). Additionally, inverse correlation of G values with Chl content has been reported as well (9,14,19,55,58). However, G has also been found to have a strong inverse correlation with Car content (21,47), which may be corroborated with the absorbance by Car within the blue-green (450–550 nm) waveband (59,60). In the present investigation, G values were distinctly lower in the HA (Anth-rich) samples compared to the LA and MA groups (Supplementary Fig. S1b), likely due to absorbance by Anth in the blue-green (400–599 nm) region (41). Hence, the influence of Anth and Car on G values makes the implementation of greenness-based or G-centric color indices unreliable especially while monitoring the health of Anth-rich plants.

Interestingly, although leaf “redness” is commonly associated with high Anth content, it was observed that the R color feature was not affected by or correlated to Anth content at all (Supplementary Figs. S1a, S2a), but maintained a strong negative correlation with SPAD Chl-meter readings. This observation can be explained by the dominant absorptive activity of Chl in the red (∼600–699 nm) waveband, where absorbance by Anth is negligible (41). Numerous studies have reported a similar correlation of R with Chl content (9,15,19,47,55,61), suggesting that reflectance within the red waveband decreases steadily with increasing Chl content. Furthermore, another study (62) demonstrated that reduction in Chl content by onset of leaf senescence triggered a sharp increase in percentage of red reflectance. Hence, such findings along with general inactivity of Car and Anth in the red waveband suggest that quantification of red reflectance or leaf redness by machine vision is a more reliable method of estimating Chl content as compared to assessing leaf greenness.

Notably, while G-centric indices such ExG, NExG, VARI, GLI, and NGBDI differentiated the HA samples from MA and LA despite similar SPAD readings (Fig. 4a, b, d, f, g), the distinction was relatively lesser in R+G-B (Fig. 4h), where R and G values are given equal weightage, and in NDPI (Fig. 4c), where G values are totally ignored. In contrast, the TREx index, wherein R is the predominant color feature with G and B values being subtracted from it (Table 1), displayed distinctly better correlation with SPAD measurements (*R*^2^ = 0.85; *n* = 320) and maintained parity across all three sample categories (Fig. 4i). This observation, along with the strong correlation of R and SPAD readings reported earlier (9,15,19,47,55,61) as well as in the present study (*R*^2^ = 0.847; *n* = 320; Supplementary Fig. S1a), indicates that redness-based indices hold the potential for representing plant health status more reliably, and hence should be explored more extensively for crop monitoring. Interestingly, analyzing additional previously-unreported redness-based color indices, such as 2*R – B, 2*R – G, 2*R + G, 2*R + G + B, and 2*R + G – B (data not shown), i.e., indices similar to TREx where R is the dominant color attribute while B and G values have been used with different mathematical functions, resulted in slightly lower correlation with SPAD values (0.74 < *R*^2^ < 0.83) compared to TREx, but performed much better than most other conventionally-used color indices. This suggests that while TREx and R indicated Chl content and plant health most reliably, alternative redness-based indices could be investigated further for implementation in different scenarios.

Nonetheless, efficacy of both TREx and R for crop monitoring was further substantiated by the SPAD prediction models using the current dataset. Amongst the different color features and indices depicted in the present study, only R, TREx, and DGCI were used for predicting SPAD values considering their better correlation with the latter (*R*^2^ > 0.7). While all three parameters provided reliable estimates of SPAD values (0.7 < *R*^2^ < 0.8, 4.7 < RMSE < 5.6), prediction was more accurate using R and TREx (Fig. 6). Hence, as SPAD values have been well-established as a benchmark for indicating plant health, both TREx and R can be extremely useful for real-time non-invasive high-throughput crop monitoring considering their strong correlation with and the ability to predict SPAD values reliably. Thus, while similar reports are abundant for green-leaved plants (56), the present study provides the first in-depth insight into the feasibility of RGB-based crop monitoring for green-leaved and anthocyanic plants in tandem.

As this study aimed at exploring plant digital color attributes from a new perspective by taking high Anth content into account, the experiments were performed under controlled conditions using excised leaves to obtain precise correlations between color features and SPAD measurements. The next step would be to investigate the utility of the proposed method with whole plants and in situ imaging to assess its practical feasibility for large-scale implementation. Specifically, exploring different lighting environments could enhance our understanding of how external light influences color features while using this approach. Additionally, while our study effectively utilized a limited number of HA samples with very low SPAD values, future research could benefit from using a larger number of samples from Anth-rich species with very low Chl content for a more detailed assessment of changes in RGB indices, especially at lower pigment levels. Investigating stressors to further reduce Chl content in Anth-rich species could offer valuable insights into this aspect as well. Future studies could also explore simultaneous assessment of plant health and nutritional value via concurrently estimating Chl and Anth contents. Enhancing prediction accuracy through more advanced data processing tools, such as deep learning, could further expand the application of this knowledge in commercial growing systems.

## 5. Conclusion

As the interest in cultivating Anth-rich leafy vegetables is growing steadily, improvement and optimization of high-throughput approaches for monitoring such plants has become imperative. In the present study, RGB images for six types of leafy vegetables possessing varying levels of Anth were analyzed simultaneously to provide an overview of variations in different digital color features and vegetation indices due to varying pigment blends. It was observed that a majority of well-established greenness-based color indices were strongly affected by high Anth content, and gave highly dissimilar outcomes for anthocyanic and green-leaved plants. Notably, increase in Anth content relative to total Chl content affected RGB-based color indices drastically until a certain threshold. However, unlike most other color features, R was not affected by leaf Anth content at all. Hence, the redness-based color index TREx was created, which was found to correlate most strongly with Chl content measured via SPAD as compared to all other color indices. By depicting the efficacy of TREx, a simple but previously-unexplored digital color index, the current study has provided novel insights into the utility of plant image analysis with focus on redness-based color attributes for green-leaved and Anth-rich plants alike. Additionally, both R and TREx were able to predict SPAD values with considerable accuracy, establishing that redness-centric data analysis is most reliable for plant health assessment. Since our analyses yielded consistent results for both green-as well as red-leaved plants belonging to four different species, our findings suggest that these colorimetric parameters have the potential to be utilized universally. In depth analyses involving additional Anth-rich crop species will further strengthen the knowledge base and broaden the applicability of this approach for assessing plant health. Expanding this research along with further validation and refinement of the method will pave the way for more widespread use in diverse agricultural settings.

## Abbreviations

*a\**: Redness-greenness (*L*a*b\** color space)
AGRI: Augmented Green-Red Index
Anth: Anthocyanin
B: Blue channel (RGB color space)
*b*: Normalized Blue channel
*b\**: Yellowness-blueness (*L*a*b\** color space)
Car: Carotenoid
Chl: Chlorophyll
DGCI: Dark Green Color Index
ExG: Excess Green
G: Green channel (RGB color space)
*g*: Normalized Green channel
GB: Greek basil
GLI: Green Leaf Index
GMR: Green-minus-Red
GPC: Green pak choi
H: Hue (HSV color space)
HA: High anthocyanin
HSV: Hue, Saturation, Value color space
*L\**: Lightness (*L*a*b\** color space)
LA: Low anthocyanin
*L*a*b\**: Lightness, Redness-greenness, Yellowness-blueness color space
MA: Medium anthocyanin
NDPI: Normalized Difference Pigment Index
NExG: Normalized Excess Green
NGBDI: Normalized Green Blue Difference Index
NGRDI: Normalized Green Red Difference Index
PB: Purple basil
R: Red channel (RGB color space)
*r*: Normalized Red channel
RGB: Red, Green, Blue color space
RPC: Red pak choi
RMSE: Root-mean-squared error
S: Saturation (HSV color space)
SK: Scarlet kale
SPAD: SPAD-502 chlorophyll meter
TREx: Two-fold Red Excess
V: Value (HSV color space)
VARI: Visible Atmospherically Resistance Index
WR: Arugula cv. ‘Wasabi Rocket’

## Acknowledgments

We thank the InFarm UK team for supplying seedlings and providing technical support, along with InFarm Crop Science team (Germany) for their support. We acknowledge all the partners (RoboScientific, Marks and Spencer, and InFarm) for their feedback and support in the project. We also thank the staff at Newcastle University for their technical, administrative and logistic support.

## Author contributions

AA: Conceptualization, Methodology, Software, Investigation, Formal analysis, Data curation, Writing-Original draft preparation, Visualization; FdJC: Resources, Methodology; VACG: Supervision, Writing-Reviewing and Editing, Project administration, Funding acquisition, Resources; TRH: Supervision, Writing-Reviewing and Editing, Project administration, Funding acquisition, Resources; NB: Supervision, Writing-Reviewing and Editing, Project administration, Funding acquisition, Resources; AP: Conceptualization, Supervision, Validation, Writing-Reviewing and Editing, Funding acquisition.

## Availability of data and materials

All data supporting the findings of this study are available within the paper and its Supplementary Information.

## Declarations

## Ethics approval and consent to participate

Not applicable.

## Consent for publication

Not applicable.

## References

1. Randhawa MA, Khan AA, Javed MS, Sajid MW. Green leafy vegetables: A health promoting source. In: Handbook of Fertility: Nutrition, Diet, Lifestyle and Reproductive Health. Elsevier Inc.; 2015. p. 205–20.

2. Yousuf B, Gul K, Wani AA, Singh P. Health benefits of anthocyanins and their encapsulation for potential use in food systems: A review. Crit Rev Food Sci Nutr. 2016;56:2223–30.

3. Gioia F Di, Tzortzakis N, Rouphael Y, Kyriacou MC, Sampaio SL, Ferreira ICFR, et al. Grown to be blue— antioxidant properties and health effects of colored vegetables. Part II: Leafy, fruit, and other vegetables. Antioxidants. 2020;9:97.

4. Carter GA, Knapp AK. Leaf optical properties in higher plants: linking spectral characteristics to stress and chlorophyll concentration. Am J Bot. 2001;88:677–84.

5. Rorie RL, Purcell LC, Mozaffari M, Karcher DE, Andy King C, Marsh MC, et al. Association of “Greenness” in corn with yield and leaf Nitrogen concentration. Agron J. 2011;103:529–35.

6. Wang Y, Hu X, Jin G, Hou Z, Ning J, Zhang Z. Rapid prediction of chlorophylls and carotenoids content in tea leaves under different levels of nitrogen application based on hyperspectral imaging. J Sci Food Agric. 2019;99:1997–2004.

7. Uddling J, Gelang-Alfredsson J, Piikki K, Pleijel H. Evaluating the relationship between leaf chlorophyll concentration and SPAD-502 chlorophyll meter readings. Photosynth Res. 2007;91:37–46.

8. Zhu J, Tremblay N, Liang Y. Comparing SPAD and atLEAF values for chlorophyll assessment in crop species. Can J Soil Sci. 2012;92:645–8.

9. Riccardi M, Mele G, Pulvento C, Lavini A, D’Andria R, Jacobsen SE. Non-destructive evaluation of chlorophyll content in quinoa and amaranth leaves by simple and multiple regression analysis of RGB image components. Photosynth Res. 2014;120:263–72.

10. Wang Y, Yang Z, Gert K, Khan HA. The impact of variable illumination on vegetation indices and evaluation of illumination correction methods on chlorophyll content estimation using UAV imagery. Plant Methods. 2023;19:51.

11. Humplík JF, Lazár D, Husičková A, Spíchal L. Automated phenotyping of plant shoots using imaging methods for analysis of plant stress responses - A review. Plant Methods. 2015;11:29.

12. Waiphara P, Bourgenot C, Compton LJ, Prashar A. Optical imaging resources for crop phenotyping and stress detection. In: Duque P, Szakonyi D, editors. Methods in Molecular Biology. New York: Humana; 2022. p. 255–65.

13. Kawashima S, Nakatani M. An algorithm for estimating chlorophyll content in leaves using a video camera. Ann Bot. 1998;81:49–54.

14. Vollmann J, Walter H, Sato T, Schweiger P. Digital image analysis and chlorophyll metering for phenotyping the effects of nodulation in soybean. Comput Electron Agric. 2011;75:190–5.

15. Agarwal A, Dutta Gupta S. Assessment of spinach seedling health status and chlorophyll content by multivariate data analysis and multiple linear regression of leaf image features. Comput Electron Agric. 2018;152:281–9.

16. Bock CH, Barbedo JGA, Del Ponte EM, Bohnenkamp D, Mahlein AK. From visual estimates to fully automated sensor-based measurements of plant disease severity: status and challenges for improving accuracy. Phytopathol Res. 2020;2:9.

17. Li D, Li C, Yao Y, Li M, Liu L. Modern imaging techniques in plant nutrition analysis: A review. Comput Electron Agric. 2020;174:105459.

18. Kim JY, Chung YS. A short review of RGB sensor applications for accessible high-throughput phenotyping. J Crop Sci Biotechnol. 2021;24:495–9.

19. Yadav SP, Ibaraki Y, Gupta SD. Estimation of the chlorophyll content of micropropagated potato plants using RGB based image analysis. Plant Cell Tissue Organ Cult. 2010;100:183–8.

20. Hu H, Zhang J, Sun X, Zhang X. Estimation of leaf chlorophyll content of rice using image color analysis. Canadian J Remote Sens. 2013;39:185–90.

21. Rigon JPG, Capuani S, Fernandes DM, Guimarães TM. A novel method for the estimation of soybean chlorophyll content using a smartphone and image analysis. Photosynthetica. 2016;54:559–66.

22. Yuan Y, Wang X, Shi M, Wang P. Performance comparison of RGB and multispectral vegetation indices based on machine learning for estimating Hopea hainanensis SPAD values under different shade conditions. Front Plant Sci. 2022;13:928953.

23. Askey BC, Dai R, Lee WS, Kim J. A noninvasive, machine learning–based method for monitoring anthocyanin accumulation in plants using digital color imaging. Appl Plant Sci. 2019;7:e11301.

24. Kim C, van Iersel MW. Image-based phenotyping to estimate anthocyanin concentrations in lettuce. Front Plant Sci. 2023;14:1155722.

25. Clemente AA, Maciel GM, Siquieroli ACS, Gallis RB de A, Luz JMQ, Sala FC, et al. Nutritional characterization based on vegetation indices to detect anthocyanins, carotenoids, and chlorophylls in mini-lettuce. Agronomy. 2023;13:1403.

26. Huang WD, Lin KH, Hsu MH, Huang MY, Yang ZW, Chao PY, et al. Eliminating interference by anthocyanin in chlorophyll estimation of sweet potato (Ipomoea batatas L.) leaves. Bot Stud. 2014;55:11.

27. Gitelson AA, Chivkunova OB, Merzlyak MN. Nondestructive estimation of anthocyanins and chlorophylls in anthocyanic leaves. Am J Bot. 2009;96:1861–8.

28. Gitelson AA, Keydan GP, Merzlyak MN. Three-band model for noninvasive estimation of chlorophyll, carotenoids, and anthocyanin contents in higher plant leaves. Geophys Res Lett. 2006;33:L11402.

29. Markwell J, Osterman JC, Mitchell JL. Calibration of the Minolta SPAD-502 leaf chlorophyll meter. Photosynth Res. 1995;46:467–72.

30. Lichtenthaler HK. Chlorophylls and carotenoids: Pigments of photosynthetic biomembranes. Methods Enzymol. 1987;148:350–82.

31. Mancinelli AL, Rabino I. Photoregulation of anthocyanin synthesis X. Dependence on photosynthesis of high irradiance response anthocyanin synthesis in Brassica oleracea leaf disks and Spirodela polyrrhiza. Plant Cell Physiol. 1984;25:1153–60.

32. Woebbecke DM, Meyer GE, Von Bargen K, Mortensen DA. Color indices for weed identification under various soil, residue, and lighting conditions. Transactions of the ASAE. 1995;38:259–69.

33. Peñuelas J, Filella I. Visible and near-infrared reflectance techniques for diagnosing plant physiological status. Trends Plant Sci. 1998;3:151–6.

34. Adamsen FJ, Pinter PJ, Barnes Robert L LaMorte EM, Wall Steven W GW. Measuring wheat senescence with a digital camera. Crop Sci. 1999;39:719–24.

35. Louhaichi M, Borman MM, Johnson DE. Spatially located platform and aerial photography for documentation of grazing impacts on wheat. Geocarto Int. 2001;16:65–70.

36. Gitelson AA, Kaufman YJ, Stark R, Rundquist D. Novel algorithms for remote estimation of vegetation fraction. Remote Sens Environ. 2002;80:76–87.

37. Karcher DE, Richardson MD. Quantifying turfgrass color using digital image analysis. Crop Sci. 2003;43:943–51.

38. Wang Y, Wang D, Zhang G, Wang J. Estimating nitrogen status of rice using the image segmentation of G-R thresholding method. Field Crops Res. 2013;149:33–9.

39. Zhang L, Zhang H, Han W, Niu Y, Chávez JL, Ma W. The mean value of gaussian distribution of excess green index: A new crop water stress indicator. Agric Water Manag. 2021;251:106866.

40. Agarwal A, de Jesus Colwell F, Bello Rodriguez J, Sommer S, Correa Galvis VA, Hill T, et al. Monitoring root rot in flat-leaf parsley via machine vision by unsupervised multivariate analysis of morphometric and spectral parameters. Eur J Plant Pathol. 2024;169:359–77.

41. Sims DA, Gamon JA. Relationships between leaf pigment content and spectral reflectance across a wide range of species, leaf structures and developmental stages. Remote Sens Environ. 2002;81:337–54.

42. Gitelson A, Merzlyak MN. Quantitative estimation of chlorophyll-a using reflectance spectra: Experiments with autumn chestnut and maple leaves. J Photochem Photobiol B. 1994;22:247–52.

43. Manetas Y. Why some leaves are anthocyanic and why most anthocyanic leaves are red? Flora. 2006;201:163–77.

44. Hughes NM, Morley CB, Smith WK. Coordination of anthocyanin decline and photosynthetic maturation in juvenile leaves of three deciduous tree species. New Phytol. 2007;175:675–85.

45. Shen J, Zou Z, Zhang X, Zhou L, Wang Y, Fang W, et al. Metabolic analyses reveal different mechanisms of leaf color change in two purple-leaf tea plant (Camellia sinensis L.) cultivars. Hortic Res. 2018;5:7.

46. Hosseini A, Zare Mehrjerdi M, Aliniaeifard S, Seif M. Photosynthetic and growth responses of green and purple basil plants under different spectral compositions. Physiol Molec Biol Plants. 2019;25:741–52.

47. Agarwal A, Dongre PK, Dutta Gupta S. Smartphone-assisted real-time estimation of chlorophyll and carotenoid concentrations and ratio using the inverse of red and green digital color features. Theor Exp Plant Physiol. 2021;33:293–302.

48. Loh FCW, Grabosky JC, Bassuk NL. Using the SPAD 502 meter to assess chlorophyll and nitrogen content of Benjamin fig and cottonwood leaves. Horttechnology. 2002;12:682–6.

49. Widjaja Putra BT, Soni P. Enhanced broadband greenness in assessing Chlorophyll a and b, Carotenoid, and Nitrogen in Robusta coffee plantations using a digital camera. Precis Agric. 2018;19:238–56.

50. Gitelson A. Towards a generic approach to remote non-invasive estimation of foliar carotenoid-to-chlorophyll ratio. J Plant Physiol. 2020;252:153227.

51. Song G, Wang Q. Developing hyperspectral indices for assessing seasonal variations in the ratio of chlorophyll to carotenoid in deciduous forests. Remote Sens. 2022;14:1324.

52. Merzlyak MN, Gitelson AA, Chivkunova OB, Rakitin VY. Non-destructive optical detection of pigment changes during leaf senescence and fruit ripening. Physiol Plant. 1999;106:135–41.

53. Merzlyak MN, Gitelson AA, Chivkunova OB, Solovchenko AE, Pogosyan SI. Application of reflectance spectroscopy for analysis of higher plant pigments. Russian J Plant Physiol. 2003;50:785–92.

54. Ku HH, Kim SH, Choi KS, Eom HY, Lee SE, Yun SG, et al. Nondestructive and rapid estimation of chlorophyll content in rye leaf using digital camera. Korean J Crop Science. 2004;49:41–5.

55. Liu Y, Hatou K, Aihara T, Kurose S, Akiyama T, Kohno Y, et al. A robust vegetation index based on different uav rgb images to estimate SPAD values of naked barley leaves. Remote Sens. 2021;13:686.

56. Kior A, Yudina L, Zolin Y, Sukhov V, Sukhova E. RGB imaging as a tool for remote sensing of characteristics of terrestrial plants: a review. Plants. 2024;13:1262.

57. Pagola M, Ortiz R, Irigoyen I, Bustince H, Barrenechea E, Aparicio-Tejo P, et al. New method to assess barley nitrogen nutrition status based on image colour analysis. Comput Electron Agric. 2009;65:213–8.

58. Jia L, Chen X, Zhang F, Buerkert A, Römheld V. Use of digital camera to assess nitrogen status of winter wheat in the Northern China Plain. J Plant Nutr. 2004;27:441–50.

59. Zang LY, Sommerburg O, Van Kuijk FJGM. Absorbance changes of carotenoids in different solvents. Free Radic Biol Med. 1997;23:1086–9.

60. Fraser PD, Pinto MES, Holloway DE, Bramley PM. Application of high-performance liquid chromatography with photodiode array detection to the metabolic profiling of plant isoprenoids. The Plant Journal. 2000;24:551–8.

61. Chen Z, Wang F, Zhang P, Ke C, Zhu Y, Cao W, et al. Skewed distribution of leaf color RGB model and application of skewed parameters in leaf color description model. Plant Methods. 2020;16:23.

62. Richardson AD, Braswell BH, Hollinger DY, Jenkins JP, Ollinger S V. Near-surface remote sensing of spatial and temporal variation in canopy phenology. Ecological Applications. 2009;19:1417–28.

