## Supplementary material for "Two-fold Red Excess (TREx): A simple and novel digital color index that enables non-invasive real-time health assessment of green-leaved as well as anthocyanin-rich crops"

Avinash Agarwal[1], Filipe de Jesus Colwell[2], Viviana Andrea Correa Galvis[2], Tom R. Hill[3], Neil Boonham[1], Ankush Prashar[1]
[1]*School of Natural and Environmental Sciences, Newcastle University, Newcastle upon Tyne, UK*
[2]*Crop Science R&D Division Infarm - Indoor Urban Farming B.V., Amsterdam, The Netherlands*
[3]*Human Nutrition and Exercise Research Centre, Population Health Science Institute, Faculty of Medical Sciences, Newcastle University, Newcastle upon Tyne, UK*

***SUPPLEMENTARY MATERIAL***

***Supplementary table***

**Table S1** Five-fold cross-validation (CV) of SPAD prediction using the Red (R) color feature.

| Dataset | Samples | | Equation | Train | | Test | |
|---|---|---|---|---|---|---|---|
| | Train | Test | | $R^2$ | RMSE | $R^2$ | RMSE |
| Full* | 320 | 320 | SPAD = 0.0009R² - 0.5106R + 74.639 | **0.791** | **4.735** | - | - |
| CV_1 | 256 | 64 | SPAD = 0.001R² - 0.5296R + 76.128 | 0.799 | 4.572 | 0.809 | 5.557 |
| CV_2 | 256 | 64 | SPAD = 0.001R² - 0.5451R + 76.052 | 0.803 | 4.67 | 0.778 | 5.349 |
| CV_3 | 256 | 64 | SPAD = 0.0009R² - 0.5131R + 74.504 | 0.784 | 4.776 | 0.825 | 4.639 |
| CV_4 | 256 | 64 | SPAD = 0.0008R² - 0.486R + 73.388 | 0.796 | 4.794 | 0.77 | 4.51 |
| CV_5 | 256 | 64 | SPAD = 0.0008R² - 0.4876R + 73.594 | 0.779 | 4.823 | 0.835 | 4.386 |
| | | | ***CV average*** | **0.792** | **4.727** | **0.803** | **4.888** |

CV_1 – CV_5, cross-validation trials with 80:20 train-test data split; $R^2$, coefficient of determination; RMSE, root-mean-squared error; $p < 0.001$. *The entire dataset was used for model creation as well as prediction.

**Table S2** Five-fold cross-validation (CV) of SPAD prediction using Two-fold Red Excess (TREx) index.

| Dataset | Samples | | Equation | Train | | Test | |
|---|---|---|---|---|---|---|---|
| | Train | Test | | $R^2$ | RMSE | $R^2$ | RMSE |
| Full* | 320 | 320 | SPAD = 0.0008TREx² - 0.3857TREx + 52.498 | **0.753** | **5.152** | - | - |
| CV_1 | 256 | 64 | SPAD = 0.0008TREx² - 0.3726TREx + 52.182 | 0.736 | 5.238 | 0.814 | 4.899 |
| CV_2 | 256 | 64 | SPAD = 0.0009TREx² - 0.3984TREx + 52.676 | 0.759 | 5.141 | 0.735 | 5.213 |
| CV_3 | 256 | 64 | SPAD = 0.0009TREx² - 0.3949TREx + 52.724 | 0.744 | 5.199 | 0.783 | 4.98 |
| CV_4 | 256 | 64 | SPAD = 0.0008TREx² - 0.3827TREx + 52.6 | 0.771 | 5.074 | 0.661 | 5.472 |
| CV_5 | 256 | 64 | SPAD = 0.0008TREx² - 0.3807TREx + 52.318 | 0.753 | 5.107 | 0.753 | 5.335 |
| | | | ***CV average*** | **0.753** | **5.152** | **0.749** | **5.18** |

CV_1 – CV_5, cross-validation trials with 80:20 train-test data split; $R^2$, coefficient of determination; RMSE, root-mean-squared error; $p < 0.001$. *The entire dataset was used for model creation as well as prediction.

**Table S3** Five-fold cross-validation (CV) of SPAD prediction using Dark Green Color Index (DGCI).

| Dataset | Samples | | Equation | Train | | Test | |
|---|---|---|---|---|---|---|---|
| | Train | Test | | $R^2$ | RMSE | $R^2$ | RMSE |
| Full* | 320 | 320 | SPAD = 115.23DGCI + 10.957 | **0.711** | **5.573** | - | - |
| CV_1 | 256 | 64 | SPAD = 112.75DGCI + 11.716 | 0.691 | 5.655 | 0.788 | 5.269 |
| CV_2 | 256 | 64 | SPAD = 115.93DGCI + 10.682 | 0.708 | 5.661 | 0.727 | 5.216 |
| CV_3 | 256 | 64 | SPAD = 114.24DGCI + 11.078 | 0.705 | 5.577 | 0.733 | 5.568 |
| CV_4 | 256 | 64 | SPAD = 117.39DGCI + 10.542 | 0.735 | 5.453 | 0.589 | 6.044 |
| CV_5 | 256 | 64 | SPAD = 115.86DGCI + 10.767 | 0.713 | 5.506 | 0.704 | 5.839 |
| | | | ***CV average*** | **0.710** | **5.570** | **0.708** | **5.587** |

CV_1 – CV_5, cross-validation trials with 80:20 train-test data split; $R^2$, coefficient of determination; RMSE, root-mean-squared error; $p < 0.001$. *The entire dataset was used for model creation as well as prediction.

*Supplementary figures*

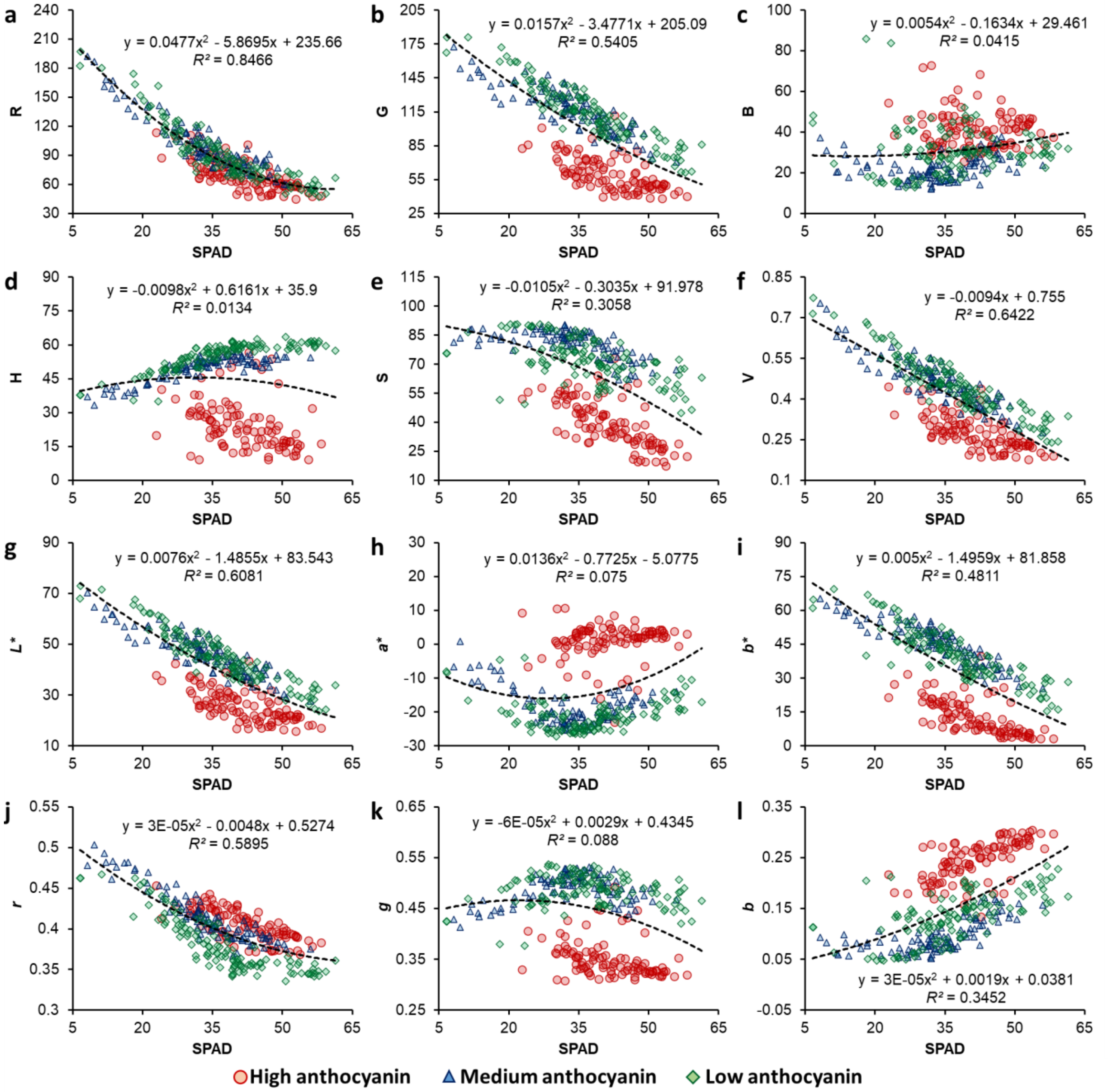


**Fig. S1** Plots of SPAD measurements with different digital color features, i.e., Red (R), Green (G), Blue (B) (a–c), Hue (H), Saturation (S), Value (V) (d–f), Lightness ($L^*$), Redness-greenness ($a^*$), Yellowness-blueness ($b^*$) (g–i), normalized Red ($r$), normalized Green ($g$), and normalized Blue ($b$) (j–l), for leaves with different levels of anthocyanin content. Coefficients of determination ($R^2$) and equations have been presented for the best-fit curve ($n$ = 320).

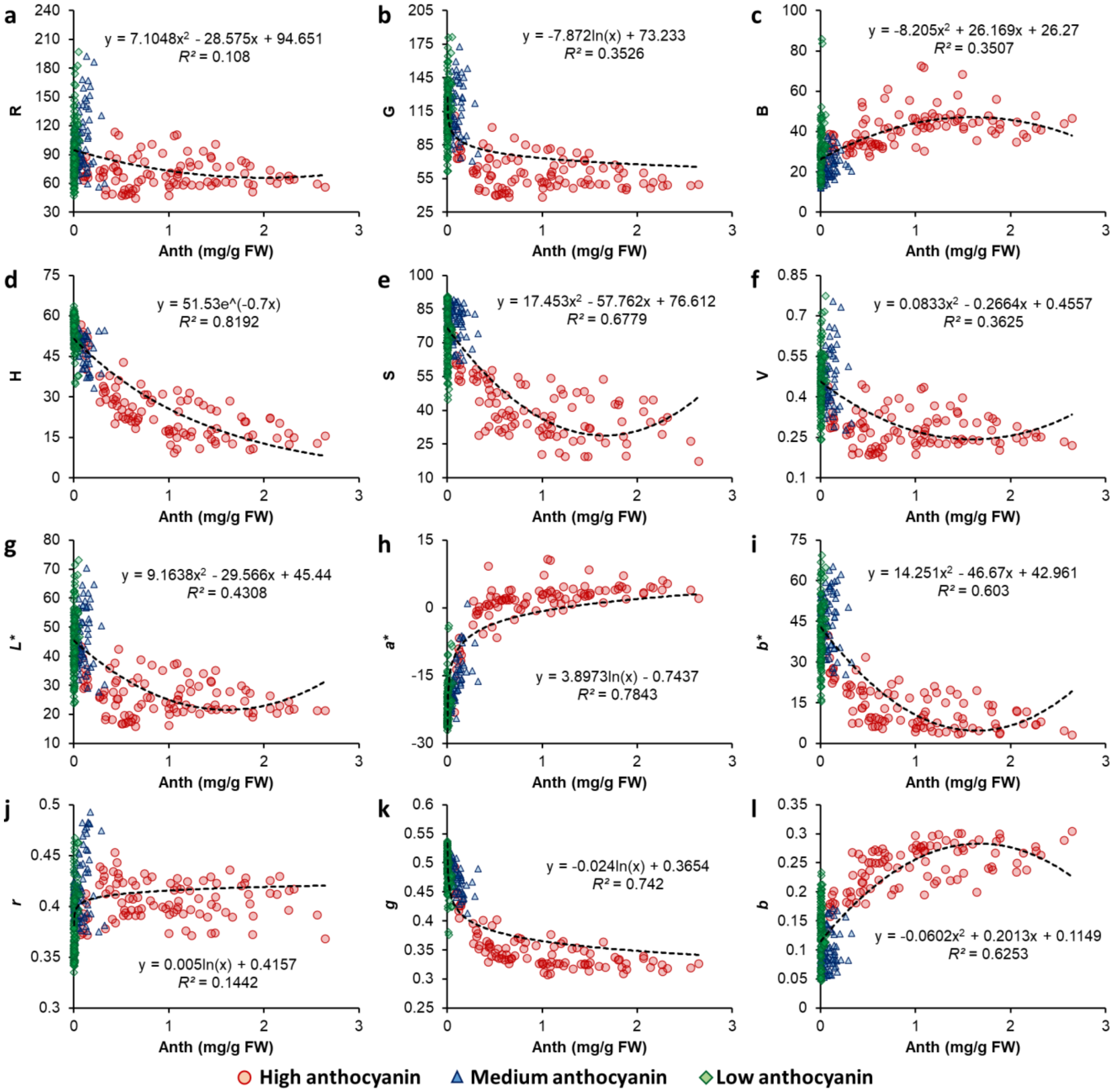


**Fig. S2** Plots of anthocyanin (Anth) content with different digital color features, i.e., Red (R), Green (G), Blue (B) (a–c), Hue (H), Saturation (S), Value (V) (d–f), Lightness (*L**), Redness-greenness (*a**), Yellowness-blueness (*b**) (g–i), normalized Red (*r*), normalized Green (*g*), and normalized Blue (*b*) (j–l), for leaves with different levels of anthocyanin content. Coefficients of determination ($R^2$) and equations have been presented for the best-fit curve ($n$ = 320).

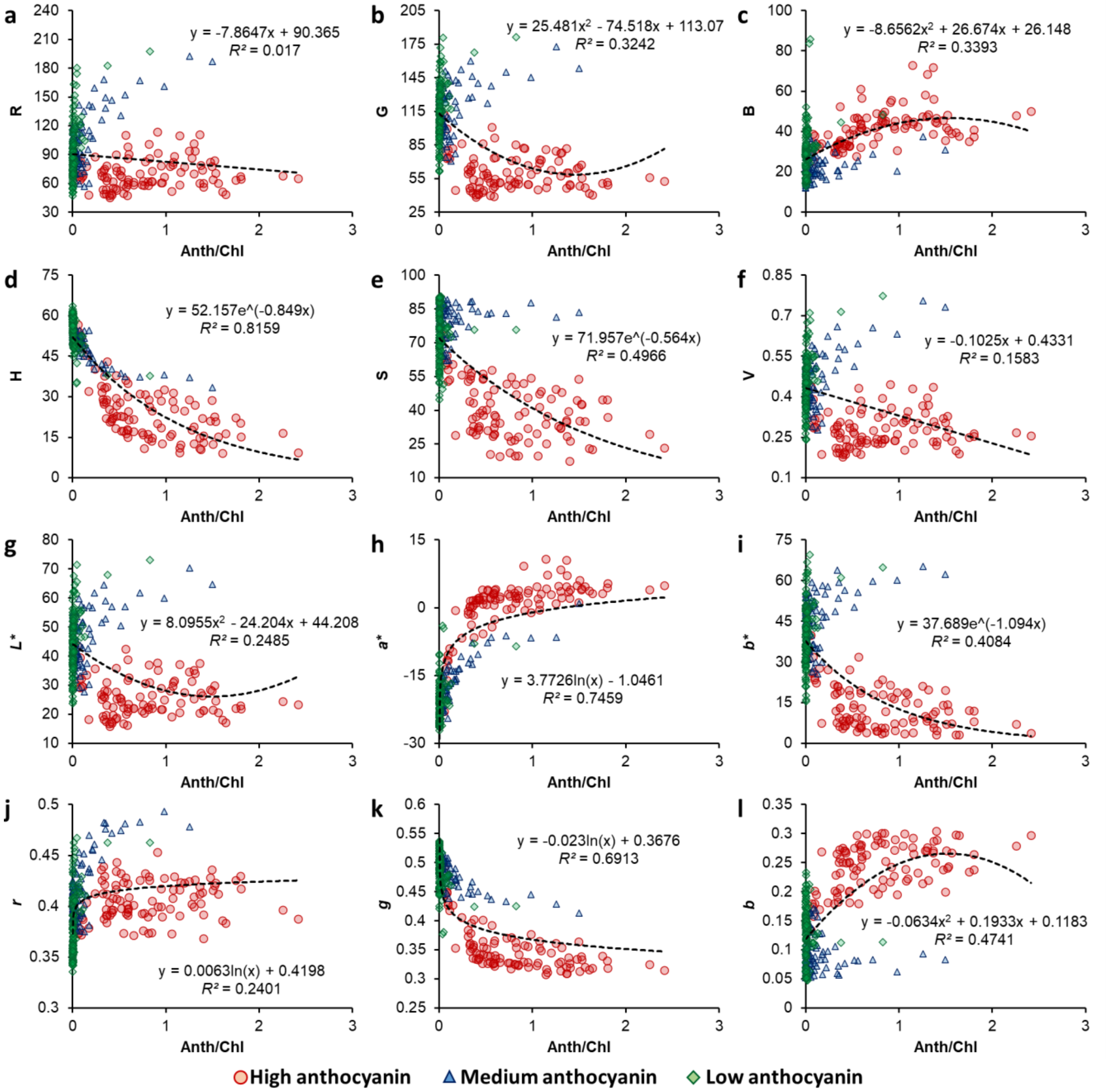


**Fig. S3** Plots of anthocyanin/chlorophyll ratio (Anth/Chl) with different digital color features, i.e., Red (R), Green (G), Blue (B) (a–c), Hue (H), Saturation (S), Value (V) (d–f), Lightness (*L**), Redness-greenness (*a**), Yellowness-blueness (*b**) (g–i), normalized Red (*r*), normalized Green (*g*), and normalized Blue (*b*) (j–l), for leaves with different levels of anthocyanin content. Coefficients of determination ($R^2$) and equations have been presented for the best-fit curve ($n$ = 320).
